# Methylome and transcriptome analysis reveals the impact of psychological stress on the skin

**DOI:** 10.1101/2025.03.14.643220

**Authors:** Bingjie Li, Ying Zou, Shenghua Tian, Laura Gonda, Andre Mahns, Tao Huang, Ludger Kolbe, Yuling Shi, Sijia Wang

## Abstract

Psychological stress has been shown to impact skin and DNA methylome, but the mechanisms are poorly understood. Here, we collected multi-omics data from 60 stressed and 60 relaxed individuals to test the hypothesis that psychological stress may impact skin via epigenetic modification and downstream altered gene expression. Each volunteer completed a series of questionnaires and assessments to measure their psychological stress levels and skin condition. Suction blister samples were collected for analyze skin cytokines, DNA methylome, and transcriptome. The dataset generated allowed for the identification of 289 differentially methylated probes(DMPs) and 10 differentially expressed genes (DEGs). Integration of methylation and expression data revealed seven functional epigenetic modules(P<0.05), which were involved in the glutamatergic synapse. In line with previous studies on the prefrontal cortex/hippocampus, we found that psychological stress also affects the glutamatergic synapse in the skin. There was no significant group difference in cytokines, while the stressed group had more severe skin darkening (P=7.36×10^-6^). This study provides important insight into the impact of psychological stress on the skin and contributes a comprehensive multi-omics dataset resource for the healthy epidermis.

## INTRODUCTION

Skin health is affected by many exposure factors (Krutmann et al., 2017), among which sun exposure (Krutmann et al., 2012), smoking (Knuutinen et al., 2002), and air pollution (Araviiskaia et al., 2019) have been extensively investigated. Recent evidence also indicates that psychological stress significantly impacts skin physiology and pathology. Chronic stress resulting from marital dissolution adversely affects skin barrier recovery, as evidenced by the elevated levels of trans-epidermal water loss (TEWL) measured 3 and 24 hours after barrier disruption (Muizzuddin et al., 2003). Delayed wound healing has been linked to higher anxiety and depression score, as assessed by the Hospital Anxiety and Depression Scale (HAD) (Cole-King and Harding, 2001). Chronic stress disrupts skin homeostasis and accelerates skin aging (Chen and Lyga, 2014, Pujos et al., 2024). In addition, psychological stress can trigger or worsen inflammatory dermatoses, such as atopic dermatitis (Arndt et al., 2008), psoriasis (Sarbu et al., 2018), contact dermatitis (Kaneko et al., 2003), acne (Jović et al., 2017), and pruritus (Steinhoff et al., 2006). However, the exact mechanisms through which psychological stress affects the skin are poorly understood. Recent studies have suggested that DNA methylation, an epigenetic mechanism that modifies gene expression, may play a role (Matosin et al., 2017). Cumulative lifetime stress has been shown to accelerate epigenetic aging, potentially driven by glucocorticoid-induced epigenetic changes (Zannas et al., 2015). The impact of high-stress response on physical signs of skin aging and whole blood epigenetic profiles has also been documented (Pattinson et al., 2016). These studies indicate that psychological stress can alter DNA methylation profile. Additionally, aberrant DNA methylation is involved in various skin conditions, with studies showing genome-wide aberrant methylation patterns and substantial transcriptomic reprogramming in UV-irradiated epidermis (Holzscheck et al., 2020), psoriasis (Zhou et al., 2016), and atopic dermatitis (Rodriguez et al., 2014). Thus, we hypothesized that psychological stress may affect skin appearance through epigenetic modifications and downstream altered gene expression. To test this hypothesis, we recruited 120 volunteers (60 stressed *versus* 60 relaxed individuals) and conducted a multi-omics analysis to compare stressed and relaxed individuals. We report that stressed individuals have an altered DNA methylome and transcriptome profile in their skin.

## RESULTS

### Participant and sample characteristics

From July 2017 to March 2018, a total of 120 healthy Chinese participants were recruited for this study, with 60 individuals in each of the stressed and relaxed groups. The psychological stress score, which was calculated from a 31-item questionnaire, was used to screen participants and ensure that the groups were appropriately classified. (**Materials and Methods**). The baseline characteristics of the study population are shown in **Table 1 and Supplementary Table S1**. Multi-omics data, including skin transcriptome (24,389 transcripts) and skin DNA methylome (736,460 CpG probes) from the lower back suction blister roofs, 36 skin cytokines (4 from the forehead rinse-off samples, 32 from the lower back suction blister fluid), and 38 skin traits (14 from skin testing, 24 from self-reported questionnaire), were collected from these participants (**Figure 1**).

**Figure 1.**
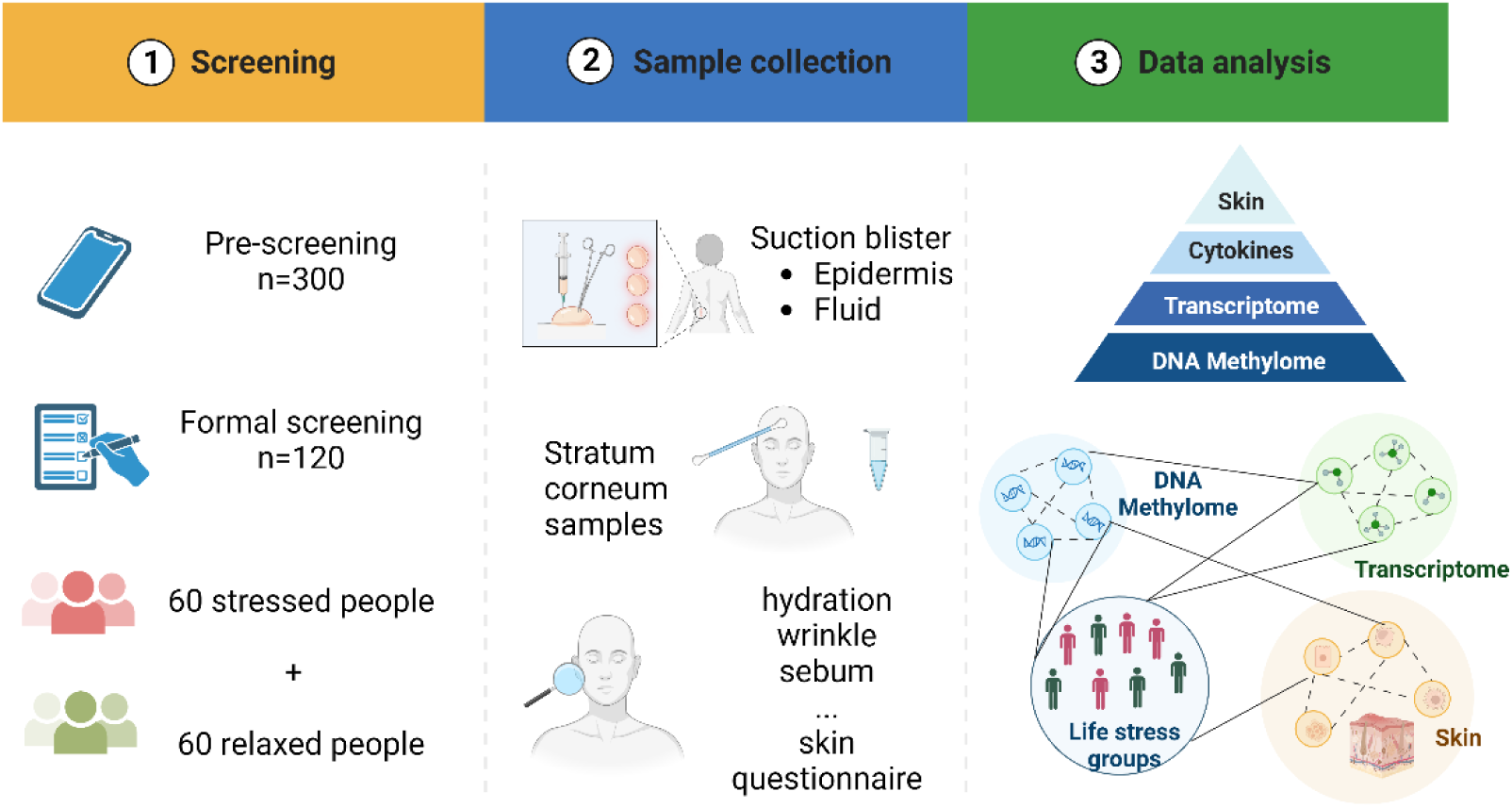
Flow diagram illustrating the study design for participant recruitment.

**Table 1.** Baseline characteristics of the study population.

|  | Relaxed group (N=60) | Stressed group (N=60) | P value<sup>1</sup> |
| --- | --- | --- | --- |
| Age (mean, sd)[min-max] | 24.18 (1.99)[21-29] | 24.15 (2.35)[19-29] | 0.93 |
| Sex (female, %) | 47 (78.33) | 45 (75.00) | 0.83 |
| BMI (mean, sd) | 20.78 (2.63) | 20.75 (4.56) | 0.97 |
| Stress score (mean, sd) | 8.4 (3.32) | 44.08 (6.39) | 4.2×10<sup>-57</sup> |

<sup>1</sup> P values were calculated using t-test for continuous variables and chi-squared tests for categorical variables. sd, standard deviation.

### Stressed individuals have an altered methylation and transcriptome profile involved in the synapse-related pathways

We analyzed genome-wide DNA methylation profiles of suction blister epidermis from 112 volunteers, comprising 56 stressed and 56 relaxed individuals. Using a Benjamini-Hochberg corrected P value of less than 0.05 and a minimum intergroup methylation difference of 1%, we identified 289 differentially methylated probes (DMPs) that corresponded to 170 annotated genes (**Figure 2a, Supplementary Table S2**). Among these DMPs, 79.58% (N=230) were significantly hypermethylated, whereas 20.42% (N=59) were hypomethylated in the stressed individuals (**Figure 2b**). Both hyper-and hypo-methylated DMPs were enriched in gene body regions (38.75%) and intergenic regions (38.75%), followed by TSS1500 regions (10.07%) (**Figure 2c**). Unsupervised hierarchical clustering analysis showed that DMPs can accurately distinguish between stressed and relaxed individuals (**Figure 2d**). The most significant DMP was located in the body of *SLC8A3* (cg01862897, Δβ=1.38%, adjusted P=2.94×10^-3^), a gene that is required for normal oligodendrocyte differentiation and myelination (Boscia et al., 2012). To identify the biological gene ontologies affected by DMPs in stressed individuals, we performed enrichment analysis using the clusterProfile package (Wu et al., 2021). Interestingly, we found 18 significantly enriched GO terms, a large proportion of which was related to lumen and synapse, such as glutamatergic synapse and neuron to neuron synapse (**Figure 2e, Supplementary Table S3**).

**Figure 2.**
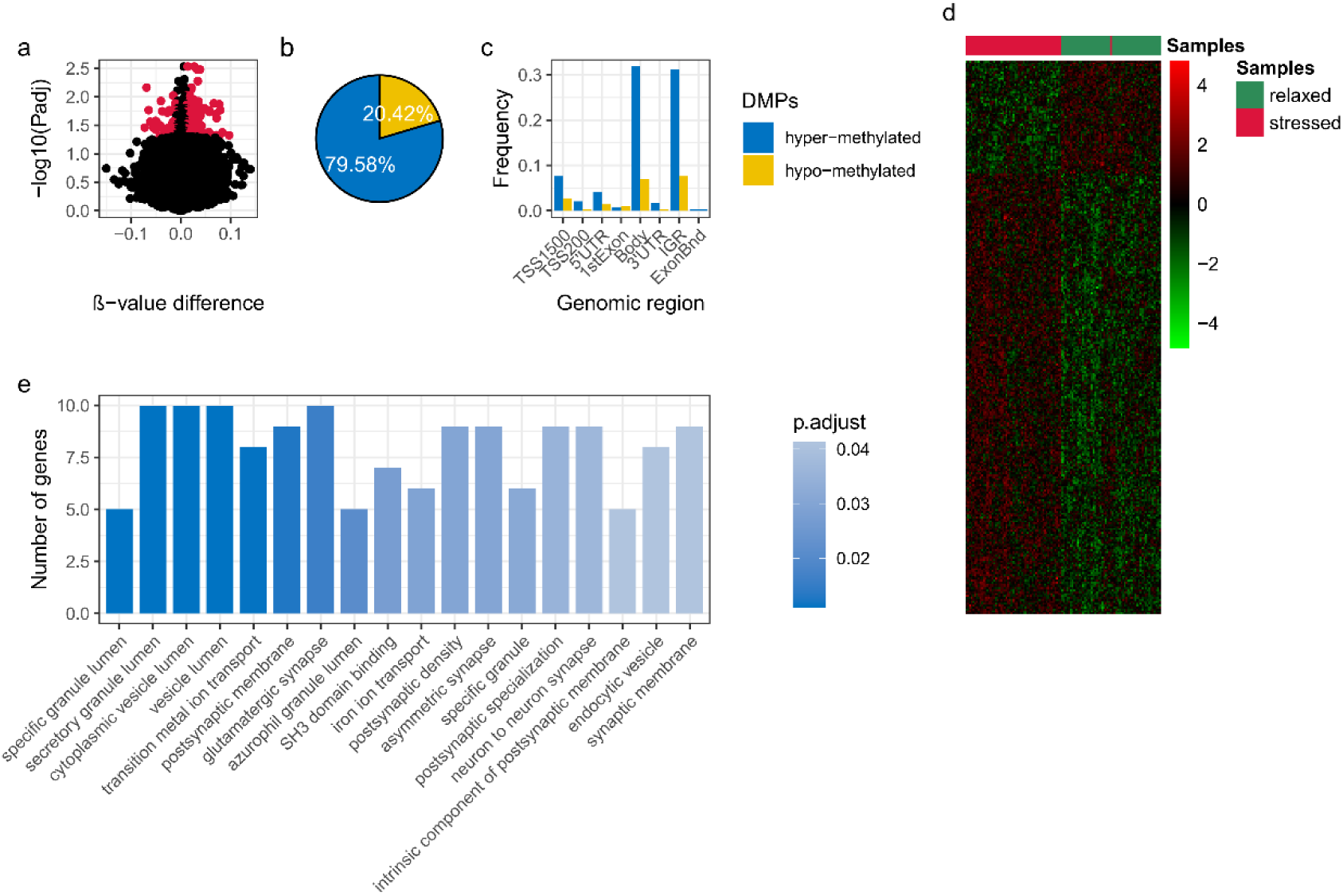
Stressed individuals have an altered methylation profile compared to relaxed individuals. a. Visualization of differential DNA methylation between the stressed (n=56) and relaxed (n=56) groups after control for age and sex. Probes with Benjamini-Hochberg adjusted p<0.05 and |Δβ| >1% were highlighted. b. Proportions of hyper-and hypo-methylated probes. c. Distribution of DMPs across genomic regions. TSS1500 (200–1500 bases upstream from the transcriptional start site, TSS), TSS200 (0–200 bases upstream from the TSS), 5’UTR (5′ untranslated region), 3’UTR (3′ untranslated region), IGR (intergenic region). d. Unsupervised hierarchical clustering of samples based on 289 DMPs. e. Significantly enriched gene ontologies based on DMPs.

To compare the gene expression patterns between stressed and relaxed individuals, we conducted RNA-seq analysis on the epidermis of 109 volunteers, comprising 53 stressed and 56 relaxed individuals. Differential expression genes (DEGs) were identified using Benjamini-Hochberg corrected P<0.05 and |log2 fold change|>1. We found that only 10 genes were differentially regulated in the stressed individuals compared to the relaxed individuals (**Figure 3, Supplementary Table S4**). Among these DEGs, 4 genes showed upregulation, while 6 genes showed downregulation in the stressed individuals.

**Figure 3.**
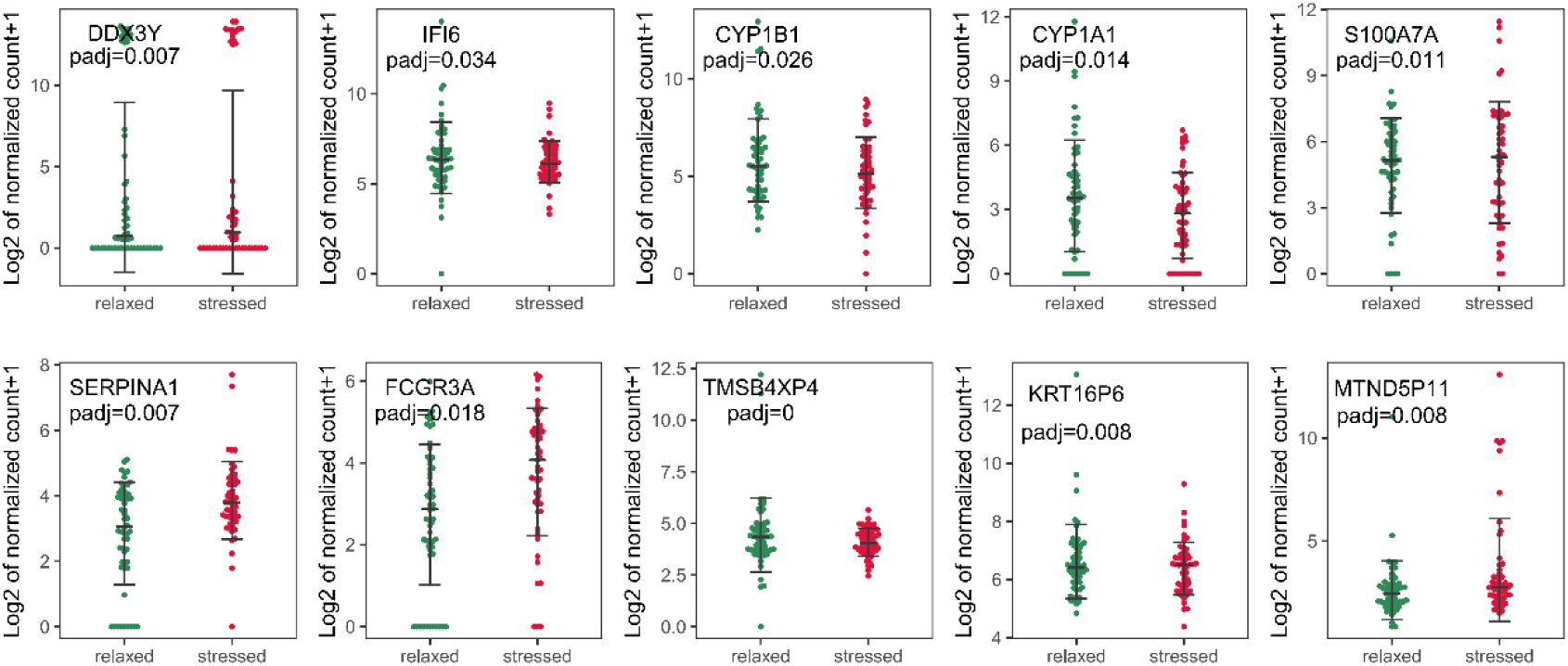
Differential expression genes between stressed and relaxed individuals. Differential expression genes (adjusted P<0.05 and |log_2_ fold change|>1) between stressed (n=53) and relaxed (n=56) groups were calculated based on the normalized read counts using DESeq2. The normalized counts were calculated by DESeq2’s median of ratios method. Vertical bars correspond to the standard deviation of the mean.

We applied a functional epigenetic modules (FEM) analysis (Jiao et al., 2014) to identify interactome hotspots of simultaneous differential methylation and gene expression associated with psychological stress. We identified seven FEM modules (P<0.05), comprising 190 unique genes (**Supplementary Table S5**). Among these genes, 56 exhibited significant differential methylated and expressed patterns, with eight genes in the *PPP3CA/EEF1G/NCK2/COPS7A/FGFR3* FEM modules showing an anti-correlation between DNA methylation and gene expression. To determine the biological significance of the FEM modules, we conducted a KEGG pathway enrichment analysis, which revealed the top two significant pathways also involved in glutamatergic synapse and axon guidance **(Figure 4, Supplementary Table S6)**.

**Figure 4.**
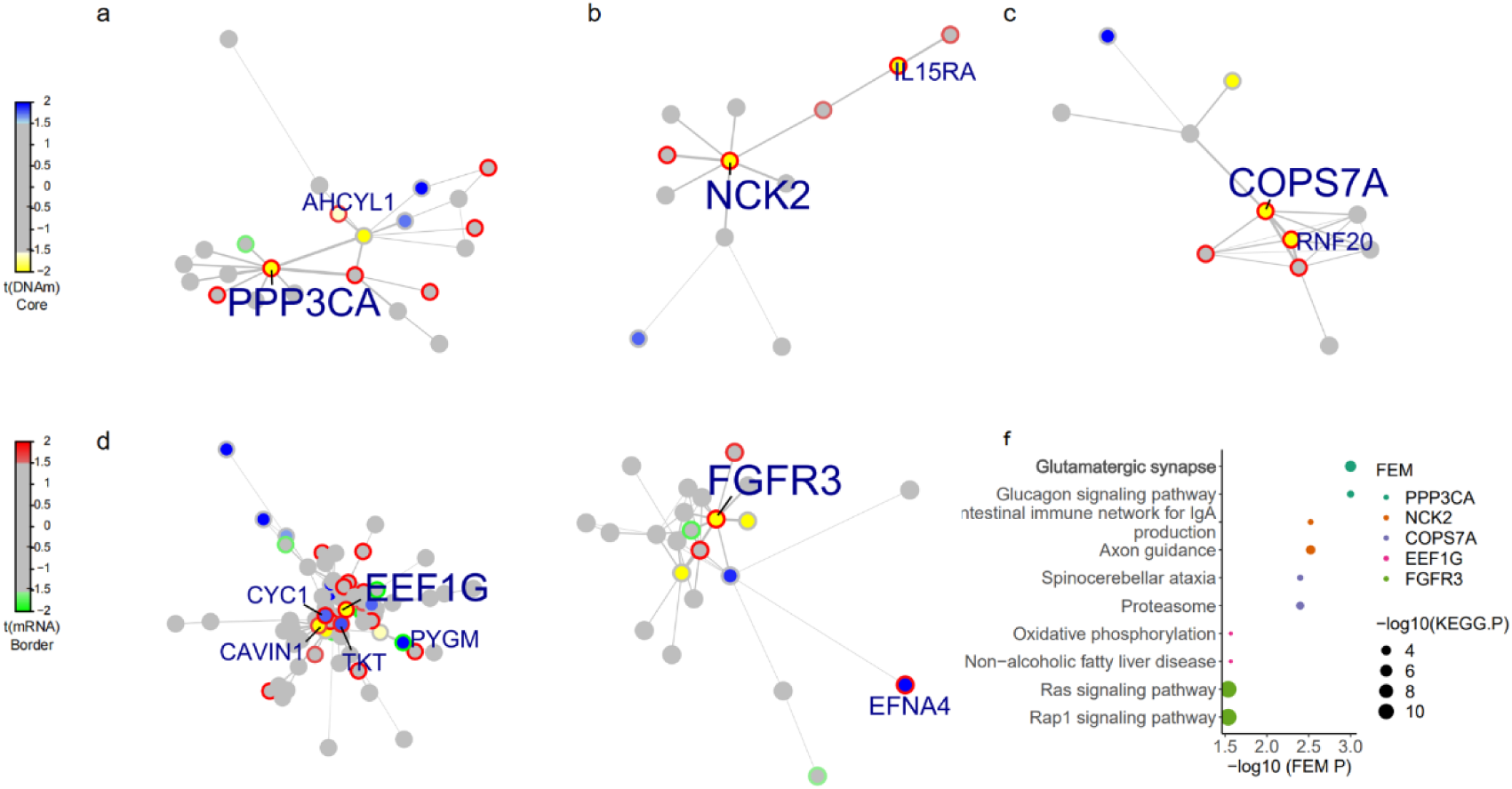
The FEM modules associated with psychological stress. a-e, Five FEMs centered around seed genes *PPP3CA, EEF1G, NCK2, COPS7A,* and *FGFR3*. Edge widths are proportional to the average statistic of the genes making up the edge. Node shades denote the differential DNA methylation statistics as indicated. Border shades denote the differential expression statistics; The labeled genes show concurrent changes in DNA methylation and gene expression levels between the stress groups. Additional genes (nodes) can be found in the Supplementary Table S5; f, significantly enriched KEGG pathway based on genes in FEM modules.

### No significant differences in the tested cytokines between stressed and relaxed groups

We compared skin phenotypes, including 36 well-known pro-and anti-inflammatory cytokines and 38 skin traits, between stressed and relaxed individuals. All cytokines were first normalized by the level of BCA protein, and the majority of cytokines exhibited a skewed distribution (**Supplementary Figure S1**). The cytokines were compared between groups using the Mann-Whitney U test, with significance levels adjusted using Bonferroni correction for multiple comparisons. We found that there was no significant difference in the tested skin cytokines collected from rinse-off samples and suction blister fluids between the two groups (**Table 2, Supplementary Figure S2**), which remained consistent across both females and males (**Supplementary Table S7**). However, as expected, the stressed group demonstrated a slight downward trend in T helper 1 (Th1) cytokines, specifically interleukin-2 (IL-2), which showed a 1.4% decrease in the average level, and interferon-gamma (IFN-γ), with a 1.5% decrease in the average level. In contrast, there was an upward trend in pro-inflammatory cytokines, such as interleukin-6 (IL-6), which exhibited a 24% increase in the average level, as well as T helper 2 (Th2) cytokines, like IL-10, which showed a 4.9% increase in the average level. These changes suggest a shift from Th1 to Th2 immune responses. Additionally, we measured skin cortisol content (collected from suction blister fluids) and found a 23% increase in the stressed group compared to the relaxed group, although this difference did not reach the significant threshold (P = 0.12).

**Table 2.** Altered skin cytokines between the stressed and relaxed groups.

| Cytokines [pg/mg]<sup>1</sup> |  | Relaxed group<br>Median (quartile)<br>(N=60) | Stressed group<br>Median (quartile)<br>(N=60) | P<br>value<sup>2</sup> |
| --- | --- | --- | --- | --- |
| Rinse off<br>samples | Lactate [ug/mg] | 274.47 (167.12-367.08) | 335.63 (175.61-396.58) | 0.43 |
|  | IL-1a | 1113.92 (826.88-1385.56) | 1200.15 (921.13-1536.92) | 0.46 |
|  | IL-1ra | 229.71 (153.85-300.29) | 256.98 (178.25-300.71) | 0.76 |
|  | IL-1ra/IL-1a | 217.95 (126.45-276.82) | 207.75 (105.14-261.79) | 0.45 |
|  | IL-8 | 157.55 (88.53-181.02) | 151.06 (93.64-172.14) | 0.88 |
| Suction<br>blister<br>fluid<br>samples | CTACK | 304.44 (210.74-412.63) | 351.38 (231.89-435.82) | 0.27 |
|  | Eotaxin | 1.23 (0.96-1.23) | 1.16 (0.97-1.23) | 0.76 |
|  | FGF.basic | 1.92 (1.63-1.86) | 1.83 (1.51-1.84) | 0.48 |
|  | G-CSF | 10.02 (8.06-11.02) | 10.62 (7.47-12.32) | 0.46 |
|  | GRO-a | 10.73 (8.27-14.38) | 10.29 (7.97-16.51) | 0.76 |
|  | HGF | 21.11 (16.57-21.94) | 19.54 (16.42-21.14) | 0.56 |
|  | IFN-a2 | 0.63 (0.51-0.63) | 0.62 (0.52-0.73) | 0.63 |
|  | IFN-g | 6.79 (5.39-6.81) | 6.18 (5.48-6.71) | 0.35 |
|  | IL-1a | 6.73 (4.14-10.37) | 6.81 (3.87-10.77) | 0.92 |
|  | IL-1b | 0.16 (0.13-0.17) | 0.16 (0.12-0.18) | 0.83 |
|  | IL-1ra | 1155.85 (978.87-1200.07) | 1224 (1073.36-1258.12) | 0.31 |
|  | IL-2 | 0.68 (0.56-0.74) | 0.64 (0.47-0.73) | 0.48 |
|  | IL-2Ra | 1.67 (1.34-1.64) | 1.52 (1.31-1.66) | 0.88 |
|  | IL-6 | 0.78 (0.27-1.37) | 0.84 (0.25-1.7) | 0.88 |
|  | IL-8 | 13.46 (6.91-15.37) | 14.56 (7.72-17.28) | 0.75 |
|  | IL-9 | 1.36 (1.14-1.83) | 1.39 (1.04-1.62) | 0.67 |
|  | IL-10 | 1.05 (0.81-1.22) | 1.17 (0.83-1.28) | 0.55 |
|  | IL-16 | 2.69 (2.12-2.72) | 2.63 (2.04-2.62) | 0.59 |
|  | IL-18 | 2.82 (2.2-2.94) | 2.49 (2.05-2.63) | 0.14 |
|  | IP-10 | 30.36 (23.42-37.25) | 29.94 (22.17-32.44) | 0.6 |
|  | MCP-1 | 41.21 (26.68-52.35) | 38.14 (27.02-64.94) | 0.93 |
|  | MIF | 495.41 (396.85-480.67) | 457.74 (390.88-494.11) | 0.91 |
|  | MIG | 12.69 (8.91-15.28) | 10.5 (8.33-13.62) | 0.25 |
|  | MIP-1a | 0.73 (0.47-1) | 0.65 (0.33-1.03) | 0.98 |
|  | MIP-1b | 5.47 (3.13-8.84) | 5.33 (3.04-10.25) | 0.77 |
|  | RANTES | 4.21 (2.67-10.93) | 4.19 (2.66-9.22) | 0.92 |
|  | SCF | 4.17 (3.32-4.54) | 3.85 (3.31-4.24) | 0.32 |
|  | SCGF-b | 2550.59 (2106.65-2754.88) | 2786.13 (2417.06-2902.86) | 0.13 |
|  | SDF-1a | 45.94 (37.93-44.58) | 45.5 (38.3-45.08) | 0.86 |
|  | TNF-a | 11.82 (8.36-14.51) | 11.85 (6.7-14.47) | 0.64 |
|  | VEGF | 10.69 (8.45-10.63) | 10.32 (8.22-10.3) | 0.5 |
|  | Cortisol | 345.73 (245.58-406.32) | 426.02 (340.11-464.18) | 0.12 |
|  | IL-1ra/IL-1a | 165.02 (92.86-260.73) | 188.77 (93.65-267.14) | 0.53 |

<sup>1</sup> Cytokines were normalized by the level of BCA protein (pg/mg).
<sup>2</sup> P values were calculated by the Mann-Whitney U test. The significant threshold after the
Bonferroni correction was $0.05/38=1.32 \times 10^{-3}$ .

Skin traits were compared using Student’s t-test and chi-squared test, with significance levels adjusted using Bonferroni correction for multiple comparisons. We found that individuals in the stressed group had more severe skin darkening than those in the relaxed group (P = 4.32×10^-5^). Moreover, the stressed group reported having less time available to perform satisfactory skincare than the relaxed group (P = 1.52×10^-10^). No other significant differences were observed in other skin traits between the two groups (**Supplementary Table S8**).

## DISCUSSION

We conducted a comprehensive multi-omics analysis to investigate the impact of psychological stress on skin, identified novel DNA methylation and expression patterns in stressed individuals. Furthermore, we have generated a valuable multi-omics dataset of healthy epidermal samples, which serves as a valuable resource for future research. Epigenetic regulation of gene expression has emerged as a crucial factor in the long-lasting impact of stress on the brain (Parade et al., 2021, Sanacora et al., 2022). Our findings revealed that psychological stress can significantly alter the skin methylome profile as well. Specifically, we identified 289 DMPs associated with psychological stress, which were significantly enriched for the lumen and synapsis-related pathways, including specific granule lumen, glutamatergic synapse, neuron to neuron synapse. Further analysis revealed a FEM module centered around *PPP3CA* that was also strongly enriched in the glutamatergic synapse pathway. These findings are in accordance with previous investigations, which have demonstrated that stress-related mediators can induce structural remodeling of dendrites and synapses within the hippocampus and prefrontal cortex (McEwen, 2012). Notably, an experimental study involving rats exposed to chronic stress for two weeks revealed significant DNA methylation changes in the prefrontal cortex, particularly within neural pathways associated with glutamatergic synapses (Wei et al., 2021). The glutamatergic system, as the vesicular neurotransmitter releaser, is known to play a crucial role in mediating the effects of psychological stress on cognition and psychopathology (Popoli et al., 2012). A recent study found that chronic stress dynamically regulates glutamatergic gene expression in the hippocampus by opening a window of epigenetic plasticity (Nasca et al., 2015). In addition, chronic stress may contribute to the pathophysiology of several psychiatric disorders through effects on the glutamatergic synapse in the prefrontal cortex (Popoli et al., 2012). Our study provides further support for the hypothesis that DNA epigenetic modifications may serve as a crucial mechanism underlying the disruption of synaptic functions in response to stress-induced perturbations. This effect is not limited to the central nervous system but extends to the peripheral nervous system (skin) as well. However, the precise mechanisms underlying these alterations are still unclear and warrant further investigation. Additionally, examination of the transcriptome data revealed 10 genes that were differentially expressed between stressed and relaxed individuals (**Supplementary Table S4**). Increased expression of *S100A7A* in hyperproliferative skin conditions such as atopic dermatitis and psoriasis has been linked to inhibited epidermal differentiation (Son et al., 2016). Upregulated expression of *S100A7A* in stressed individuals (log2 fold change = 1.599, adjusted P = 0.01) may supports previous evidence that psychological stress can trigger or exacerbate psoriasis (Xhaja et al., 2014).

In previous studies, exposure to chronic stress during academic examinations has been associated with elevated levels of plasma IL-1β, IL6, and IL10, along with reduced production of plasma IL-2 and IFN-γ (Kang et al., 2001, Paik et al., 2000). Alzheimer’s caregivers reported significantly higher levels of distress and increased IL-6 concentrations (Kiecolt-Glaser et al., 2003, Lutgendorf et al., 1999). Additionally, burnout, characterized as a stress-induced work-related syndrome, has been found to correlate with an increase in the production of the anti-inflammatory cytokine IL-10 by monocytes (Mommersteeg et al., 2006). However, in our study, we did not observe any significant changes for cytokines in the skin. There are several possible explanations for this finding. First, it is possible that unstimulated chronic psychological stress experienced by our healthy individuals may have less effect on skin cytokines in this observational study; Second, although we collected a large number of samples, detecting minor effects may require a larger sample size to provide sufficient statistical power; Third, mast cells are known to regulate neurogenic inflammation and are a source of skin cytokines during stress responses (Arck et al., 2006). The cytokines we measured in the epidermal tissue fluid may not fully represent the whole picture of the mast cells. In future studies, we recommend using whole skin samples for cytokine detection to obtain a more comprehensive understanding of the effects of stress on skin cytokines. Although the intergroup differences in cytokine levels were not statistically significant, the observed trends in their variation were consistent with prior research, suggesting that chronic psychological stress may induce a shift from Th1 to Th2 immune responses. Our analyses showed that stressed individuals tend to have darker skin compared to relaxed individuals, which is in line with a previous study concluding that fatigue induces tired-looking and dull skin in Chinese women (Flament et al., 2019). Skin darkening was assessed using a self-reported questionnaire; further studies should use more objective detection methods, such as skin code reader, for validation. The generalizability of this finding to other populations beyond the Chinese cohort requires further investigation.

Moreover, we have generated a valuable multi-omics dataset. To minimize the confounding effects of cell heterogeneity, we analyzed the DNA methylome and transcriptome specifically in the epidermal samples. Similar to previous eQTM studies in other tissues or cell lines (Gutierrez-Arcelus et al., 2015, Kim et al., 2020, Sharma et al., 2020), this dataset provides a valuable resource for performing eQTM analysis in the healthy epidermis.

Our study has some limitations. Firstly, the study’s cross-sectional design precludes the assessment of causal relationships and the dynamic effects of stress on the skin over time. A longitudinal study approach would provide more insights into the temporal changes in the examined parameters. Additionally, the relatively small sample size may have limited the ability to detect subtle effects, particularly on skin cytokine levels. Further validation of the findings in a larger, more diverse cohort, or using other methodological approaches (qRT-PCR) would be necessary to confirm the generalizability of the results. Finally, the study’s observational nature makes it challenging to fully differentiate the impacts of stress from other potential confounding lifestyle factors, such as sleep quality, pregnancy history, and dietary habits. Addressing these limitations through future longitudinal and controlled studies would strengthen the conclusions drawn from this study.

In conclusion, our study reveals distinct DNA methylation and expression signatures that define the epidermal responses to psychological stress. The findings not only enhance our understanding of the impact of stress on the epidermis but also provide a valuable dataset resource for future investigations.

## MATERIALS & METHODS

### Study participants

Between July 2017 and March 2018, we enrolled 300 healthy Han Chinese individuals from Shanghai for our study. Participants were selected based on their responses to a pre-screening psychological stress questionnaire **(Figure 1)**. Inclusion and exclusion criteria (**Supplementary data S1**) included a history of abnormal wound healing, any clinically diagnosed severe skin conditions and abnormal skin at the test areas, a history of HIV, infectious hepatitis, diabetes, cancer, rheumatic disease, cardiovascular disease, thyroid dysfunction, or asthma. We also excluded participants that taking medications for anti-phlogistic agents or analgetic, suppress immune defenses or antihistamines, or who had used blood coagulation-related drugs within the last 2 weeks, or oral or topical retinoid within the last 2 months, as well as smokers and individuals who were pregnant or breastfeeding.

To select the most extreme participants, all participants completed a detailed psychological stress questionnaire. The psychological stress questionnaire consists of 31 items (**Supplementary data S2**). Stress score is derived from four modules (Equation 1). Module A indicates the stress index and includes 12 positive items (Q1, Q3, Q5, Q6, Q8, Q12, Q14, Q15, Q22, Q26, Q29, Q30) and 4 negative items (Q10, Q16, Q23, Q25). Module B represents the relaxation index and comprises 8 positive items (Q2, Q4, Q7, Q9, Q11, Q17, Q18, Q28). Module C reflects enhanced stress and consists of 6 positive items (Q19, Q20, Q21, Q24, Q27, Q31) and a negative item (Q13). A constant of 16 is also included. Positive items are scored as 2 for “often”, 1 for “sometimes”, and 0 for “rarely”, while negative items are scored as 0 for “often”, 1 for “sometimes”, and 2 for “rarely”. The scores are calculated as follows: Score A is the total score from Module A, B is the total score from Module B, and Score C is the minimum of the sum of items scoring 2 in Module A and the sum of items scoring 2 in Module C.

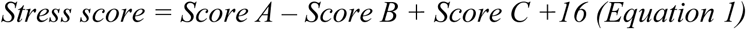

To validate the reliability of our psychological stress questionnaire, we collected both the PSS-10 questionnaire and our psychological stress questionnaire from 32 volunteers, calculating the stress scores for both instruments. The stress score obtained from the two questionnaires showed a strong correlation (r=0.75). Additionally, individuals classified as stressed and relaxed group exhibited significant differences in PSS-10 scores (P = 9.6×10^-6^). These findings support the reliability of our psychological stress questionnaire in assessing psychological stress levels (**Supplementary Figure S3**).

We calculated the stress score for each participant based on the questionnaire data and selected the top 60 stressed individuals and top 60 relaxed individuals for the next stage of the study. At the time of sample collection and skin trait measurement (conducted 5 to 10 days later), participants completed a brief stress questionnaire to confirm the continued presence of between-group differences in perceived stress levels (**Supplementary Table S9**). The age range of these participants was 19 to 29 years, with 92 women [76.67%] and 28 men [23.33%]. This study was approved by the Ethical Committee of Fudan University, Shanghai, China, and all participants provided written informed consent.

### Skin trait measurement

Prior to the measurements, participants were told to (i) not to use any kind of skincare products for the last 5 days, and (ii) not to wash face, not to perform sweat-inducing sports, and not to use a sauna or have a bath for the last 24 hours. All volunteers reported that the sampling site (lower back) had not been exposed to intense sun exposure within the 28 days prior to the sample collection. We asked participants to wait for about 30 minutes to adapt to the room environment (25℃, 50%) before the measurement. Skin physiological traits were measured using a self-assessment questionnaire and a skin code reader (Beiersdorf, Hamburg, Germany).

### Sample collection

Skin suction blisters were formed using a low-pressure device as previously described (Südel et al., 2003), by mounting one sterile 3-hole (3-mm diameter per hole) skin suction chamber onto the lower back of each participant. Negative pressure (180-240 mmHg) was applied over 1.5 to 2 hours, until blister formation was complete. Standard surgical instruments were used to collect suction blister samples (roof and fluid). Rinse-off samples (skin surface protein) were collected by repeatedly scrubbing the forehead of volunteers with a sterile cotton tip that soaked in a buffer. All samples were taken and immediately stored at-80℃.

### Cytokine

Skin cytokines levels were determined for both rinse-off and suction blister samples. For rinse-off samples, lactate level was measured by Lactate Reagent (Pointe Scientific, # 7596-50) and Lactate Standard Solution (Pointe Scientific, #7596-STD). IL-α, IL-1ra, IL-8, and BCA were measured by assay kit (R&D System, #DLA50; R&D System, # DRA00B; BD Biosciences, #550999; Thermo Scientific, #23227). For suction blister fluid, cytokine levels were measured by multiplexing with a Bio-Plex 200 System with a Bio-Plex Pro Human Cytokine 48-Plex Panel (#12007283), according to the manufacturer’s instructions. BCA was determined by the BCA kit (Thermo Scientific, #23227). BCA protein level was used for normalization. Cytokines were compared between stressed and relaxed individuals using Mann-Whitney U-test.

### Transcriptome

Suction blister roofs (epidermis) were used to isolate total RNA through RNeasy Fibrous Tissue Mini Kit (Qiagen, Hilden, Germany). Single-end sequencing was performed at 75 base pairs (bps) on the Illumina NextSeq500 system, resulting in a final sequencing depth of approximately 100 million reads per sample. RNA reads with low quality and adaptors were removed. Clean reads were aligned to the GENCODE human genome assembly GRCh38 (hg38) using Bowtie2 (Langmead and Salzberg, 2012). Read counts and FPKM values were calculated using RSEM(Li and Dewey, 2011). Differential gene expression analysis was calculated using DESeq2 (Love et al., 2014), adjusting for age, gender, and top 2 hidden batch effect calculated by RUVseq (Risso et al., 2014). Genes were considered significantly differentially expressed with |log2 fold change|>1 and adjusted P<0.05 after multiple testing adjustments by the Benjamini-Hochberg method.

### DNA methylome

Suction blister roofs (epidermis) were used to isolate DNA through DNeasy Blood & Tissue Kit (Qiagen, Hilden, Germany). Methylation profiling was performed using Infinium MethylationEPIC arrays (Illumina). Methylation data were processed using the R package ChAMP (Tian et al., 2017) pipeline. Probes were filtered out if the detection P value of >0.01 or if 5% of probes failed. Non-CpG probes, SNP-related probes, multi-hit probes, and probes located in chromosome X and Y were also removed. Finally, 736,460 probes and 112 samples (56 stressed individuals and 56 relaxed individuals) remained after quality control. Intra-array and probe design variation were corrected for using beta-mixture quantile dilatation (BMIQ (Teschendorff et al., 2013)). Inter-array batch effects were corrected for using ComBat (Johnson et al., 2007). Differentially methylated probes analysis was performed using limma (Ritchie et al., 2015) together with age and sex as covariates. Statistical significance was set at |Δβ|>1% and P<0.05 after multiple testing adjustments by the Benjamini-Hochberg procedure.

### Statistical analyses

Skin traits were compared between the stressed and relaxed group using T-tests for continuous variables and chi-squared tests for categorical variables. Cytokines were compared between the stressed and relaxed groups using the Mann-Whitney U test. Genes annotated by DMPs were used to do the enrichment analysis using clusterProfiler package (Wu et al., 2021). The integrative analysis of DNA methylation and gene expression data was performed by the FEM algorithm (Jiao et al., 2014) to identify biological modules and pathways that are related to psychological stress. FEM integrates information from DNA methylation, gene expression, and protein-protein interaction (PPI) networks, rather than just filtering DEGs or DEPs using a specific cutoff, which can provide a more comprehensive understanding of the underlying biological mechanisms.

## DATA AVAILABILITY STATEMENT

All of the DNA methylome data, transcriptome data and cytokine data have been deposited in National Omics Data Encyclopedia (http://www.biosino.org/node/) under accession number OEP003724. Additional data supporting the findings of this study are available from the corresponding author upon reasonable request.

## ACKNOWLEDGMENTS

This work was supported by the Strategic Priority Research Program (Grant No. XDB38020400), Shanghai Municipal Science and Technology Major Project (Grant No.2017SHZDZX01) to SW.

## AUTHOR CONTRIBUTIONS STATEMENT(CRediT)

Bingjie Li: Data Curation, Formal Analysis, Investigation, Visualization, Writing - Original Draft Preparation, Writing - Review and Editing

Ying Zou: Supervision, Writing - Review and Editing, Project Administration Shenghua Tian: Formal Analysis, Writing - Review and Editing

Laura Gonda: Conceptualization, Data Curation Andre Mahns: Conceptualization, Data Curation Tao Huang: Writing - Review and Editing

Ludger Kolbe: Conceptualization, Writing - Review and Editing

Yuling Shi: Supervision, Writing - Review and Editing, Project Administration

Sijia Wang: Supervision, Conceptualization, Funding Acquisition, Writing - Review and Editing, Resources, Project Administration

